# Native Mass Spectrometry Reveals the Conformational Diversity of the UVR8 Photoreceptor

**DOI:** 10.1101/371658

**Authors:** Inês S. Camacho, Alina Theisen, Linus O. Johannissen, L. Aranzazú Díaz-Ramos, John M. Christie, Gareth I. Jenkins, Bruno Bellina, Perdita Barran, Alex R. Jones

**Author notes:** **Materials & Correspondence**, To whom correspondence and material requests should be addressed.

## Abstract

UVR8 is a plant photoreceptor protein that regulates photomorphogenic and protective responses to UV light. The inactive, homodimeric state absorbs UV-B light resulting in dissociation into monomers, which are considered to be the active state and comprise a β-propeller core domain and intrinsically disordered N- and C-terminal tails. The C-terminus is required for functional binding to signalling partner COP1. To date, however, structural studies have only been conducted with the core domain where the terminal tails have been truncated. Here, we report structural investigations of full-length UVR8 using native ion mobility mass spectrometry adapted for photo-activation. We show that, whilst truncated UVR8 photo-converts from a single conformation of dimers to a single monomer conformation, the full-length protein exist in numerous conformational families. The full-length dimer adopts both a compact state and an extended state where the C-terminus is primed for activation. In the monomer the extended C-terminus destabilises the core domain to produce highly extended yet stable conformations, which we propose are the fully active states that bind COP1. Our results reveal the conformational diversity of full-length UVR8. We also demonstrate the potential power of native mass spectrometry to probe functionally important structural dynamics of photoreceptor proteins throughout nature.

**TOC Graphic:** 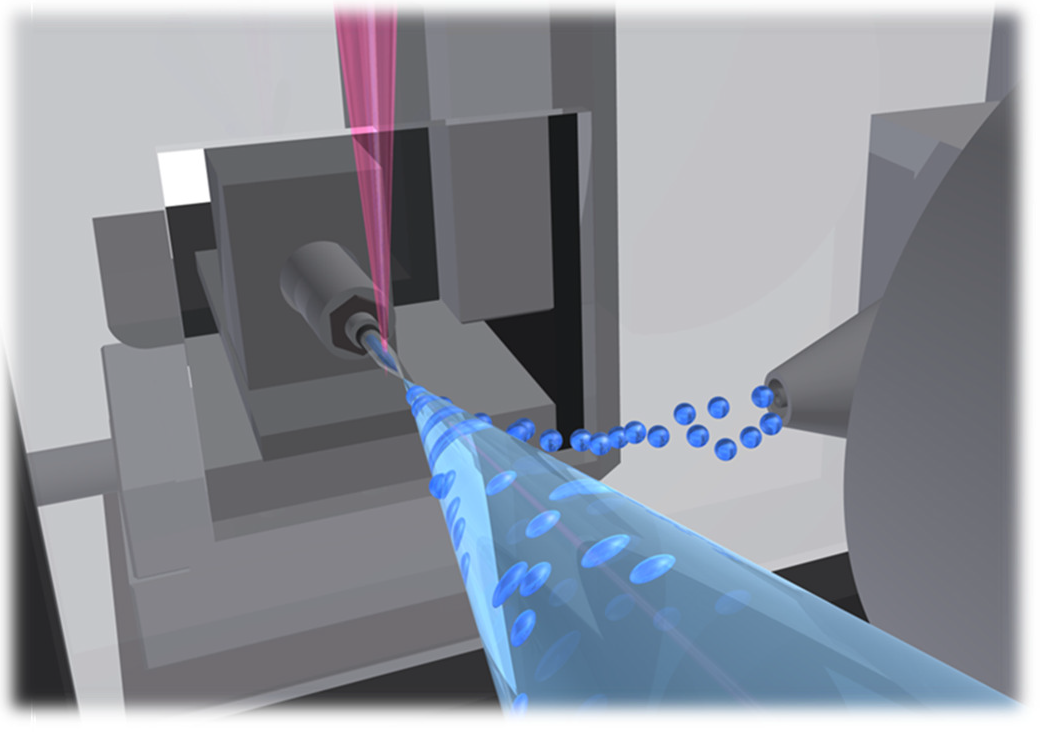

## Introduction

The functional importance of disorder in proteins is transforming the structure-function paradigm. Intrinsically disordered regions (IDRs) of proteins have been found to play a particularly important role in cell signalling pathways, where the lack of structural order provides accessible and often versatile sites for binding to signalling partners and for post-translational modification.^1^ Consistent with this, it is becoming increasingly apparent that IDRs facilitate signal propagation following activation of photoreceptor proteins by environmental light cues.^e.g., 2-6^ The plant photoreceptor, UV RESISTANCE LOCUS8 (UVR8),^7,8^ serves as an illustrative example. It comprises a structurally well-defined seven blade β-propeller core domain^9,10^ alongside what are thought to be conformationally flexible C- and N-terminal tails (Figures 1a&b).^9^ In the dark UVR8 adopts a homodimeric structure, which is stabilised by cross-dimer salt bridges (Figure 1b). Absorption of UV-B light (280-315 nm) by chromophores made up of clusters of tryptophan residues at the dimer interface (Figure 1c) results in disruption of these salt bridges, which in turn leads to dissociation into monomers^9-11^ and accumulation of UVR8 in the nucleus.^12^ Although it is widely accepted that monomerisation is necessary for UVR8 activation, it is now clear that the C-terminal tail is vital for subsequent signal transduction and for regulation of UVR8 function.^5,11,13^

**Figure 1.**
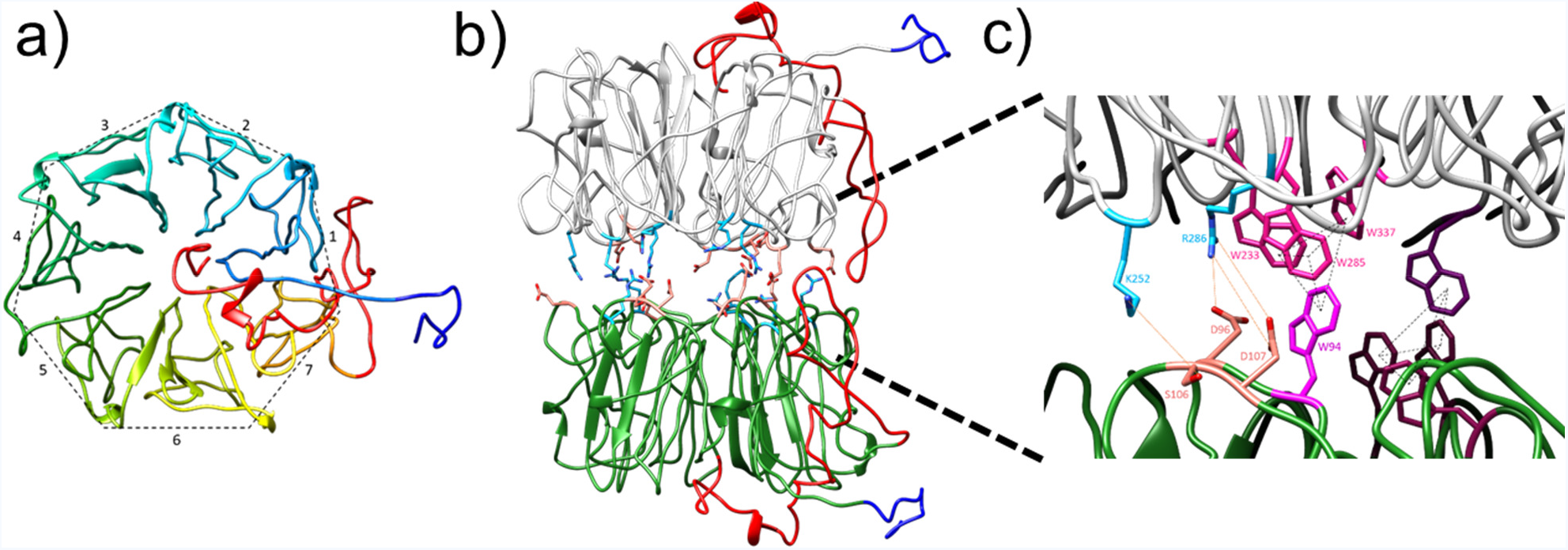
Structural features of the UVR8 photoreceptor. We have modelled in the conformationally-disordered C- (red) and N- (dark blue) terminal tails onto the published^9^ structure of the UVR8 core domain (see Supporting Information and Experimental Section for details). **a)** ‘Top’ view of a UVR8 monomer, showing the WD40, β-propeller structure of the core domain and the relative positions of the C- and N-terminal tails. **b)** ‘Side’ view of the UVR8 homodimer, which is stabilised by cross-dimer salt bridges (positive residues – light blue; negative residues – peach). **c)** The UVR8 dimer contains two chromophores (pink / purple), each of which comprises a pyramid of tryptophan residues: a triad from one monomer and a forth from the other. Each pyramid neighbours residues involved in key cross-dimer salt bridges (shown for the pink chromophore).

In the nucleus of the cell, photoactivated UVR8 monomer binds to CONSTITUTIVELY PHOTOMORPHOGENIC1 (COP1), the central regulator of light signalling in plants.^11,14^ Together they regulate over 100 genes involved in UV-B acclimation (e.g., biosynthesis of UV-absorbing flavonoids) and photomorphogenic responses such as suppression of hypocotyl growth.^14^ The C-terminus contains a highly conserved sequence of 27 amino acids (residues 397-423 of 440) that is known to be required for not only functional binding to COP1, but is also necessary and sufficient for binding to the REPRESSOR OF UV-B PHOTOMORPHOGENESIS proteins, RUP1 and RUP2, which are negative regulators of UVR8.^5,15^ Indeed, expression of just the final 44 amino acids of the C-terminus (residues 397-440) in plants is sufficient to trigger expression of the transcription factor, ELONGATED HYPOCOTYL5 (HY5),^15^ which mediates many of the responses triggered by UVR8.^13^ Despite these data supporting a functional role for the C-terminal IDR, little is known about the extent of disorder of the C-terminus or about its structural dynamics and how they relate to the active state of UVR8. Crystal structures have been solved for the UVR8 core domain (residues 12-381, UVR8^12-381^)^9,10,16^ and have yielded significant mechanistic information regarding light-induced monomerisation. In order for the protein to crystallise, however, it was necessary to truncate the N- and C-terminal tails, both implying structural disorder in these regions of UVR8 and precluding their detailed study. Here, we have acquired data of both UVR8^12-381^ and full-length UVR8 using novel native ion mobility mass spectrometry instrumentation that incorporates illumination of the sample solution at the ion source. These data not only enable monitoring of the UV-B-induced dissociation of the UVR8 dimer but also reveal the conformational diversity of the full-length protein. We believe this method will prove to be widely applicable and powerful technique with which to investigate both photoreceptor activation mechanisms and their signalling conformations.

## Results

### A simple optical assembly enables the investigation of photoreceptor structural dynamics by native mass spectrometry

Native mass spectrometry was used to investigate the photoactivation of UVR8 using a simple, versatile and modular illumination assembly (Figure 2). A high-power LED (25 mW, 350 mA, Thorlabs) emitting at a peak wavelength of 280 nm was mounted above the ion source. The output was focussed onto the nano-electrospray ionisation (nESI) tip of the ion source using convex lenses allowing irradiation of the sample solution prior to desolvation and before it enters the mass spectrometer. This illumination assembly is compatible with a range of commercial spectrometers, and has been used with spectrometers from 3 different manufacturers in this study (see Methods section). All known photoreceptor proteins are activated with wavelengths corresponding to the solar emission spectrum that reaches the earth’s surface. This range (near UV – visible – near IR) is covered by inexpensive and readily available LEDs similar to the one used here. In principle, therefore, the illumination assembly can be straightforwardly adapted for investigation of the structural changes that follow photoactivation of all light-responsive proteins – and indeed many other photochemical reactions.

**Figure 2.**
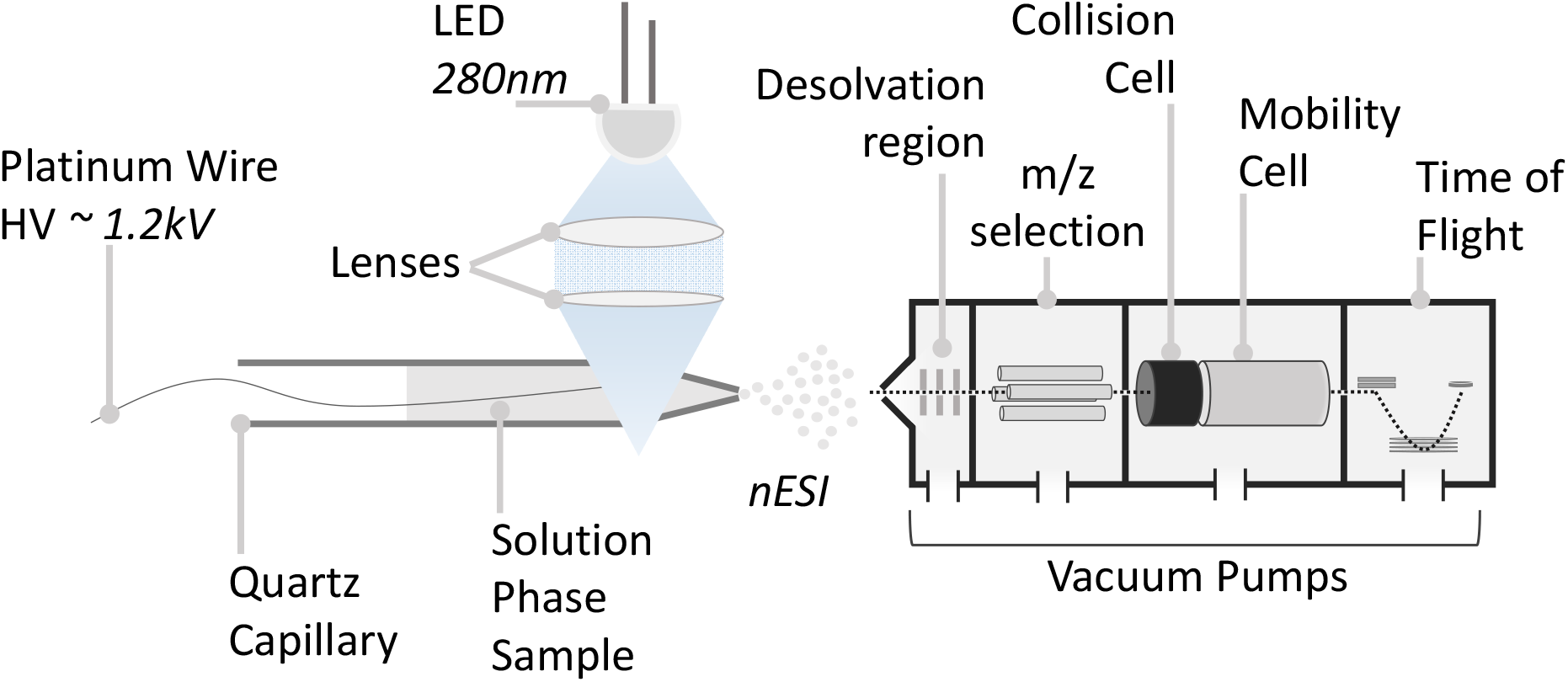
Photo-ion mobility mass spectrometry. Schematic of the experimental setup. The liquid phase sample is loaded in a quartz capillary (10 μl). Using a lens assembly the light emitted from a LED is focused at the end of the capillary. High voltage is applied to the solution resulting in the nebulisation of the sample and its transfer to the gas phase by nESI. The generated ions are then transferred into an ion mobility mass spectrometer where both collision cross section (CCS) and molecular weight (m/z) data are obtained.

### The UVR8 core domain exists in a single conformational family

Crystal structures have been solved for UVR8^12-381^ where the N- and C-terminal tails were truncated from the full-length protein.^9,10,16^ To investigate the conformational flexibility of UVR8 in the absence of its IDRs we acquired the native mass spectrum of the UVR8^12-381^ dimer in the absence of UV-B light (Figure 3a). These data reveal a narrow charge state distribution where z= 15+ to 18+, consistent with the UVR8^12-381^ dimer existing in a single conformational family. This conclusion is supported by ion mobility drift time (DT) data acquired in both helium (Figure 3b) and nitrogen (Figure S1a). A global collision cross section, ^DT^CCS_He_, of 4390 ± 3 Å^2^ was obtained from these data, and is in good agreement with the theoretical value, ^Th^CCS_He_, of 4566 ± 43 Å^2^ from molecular dynamics (MD) simulations (Figure S2a) of the energy minimised, published^9^ structure (Figure 3b inset). From these data we conclude that the UVR8^12-381^ dimer is structurally ordered.

**Figure 3.**
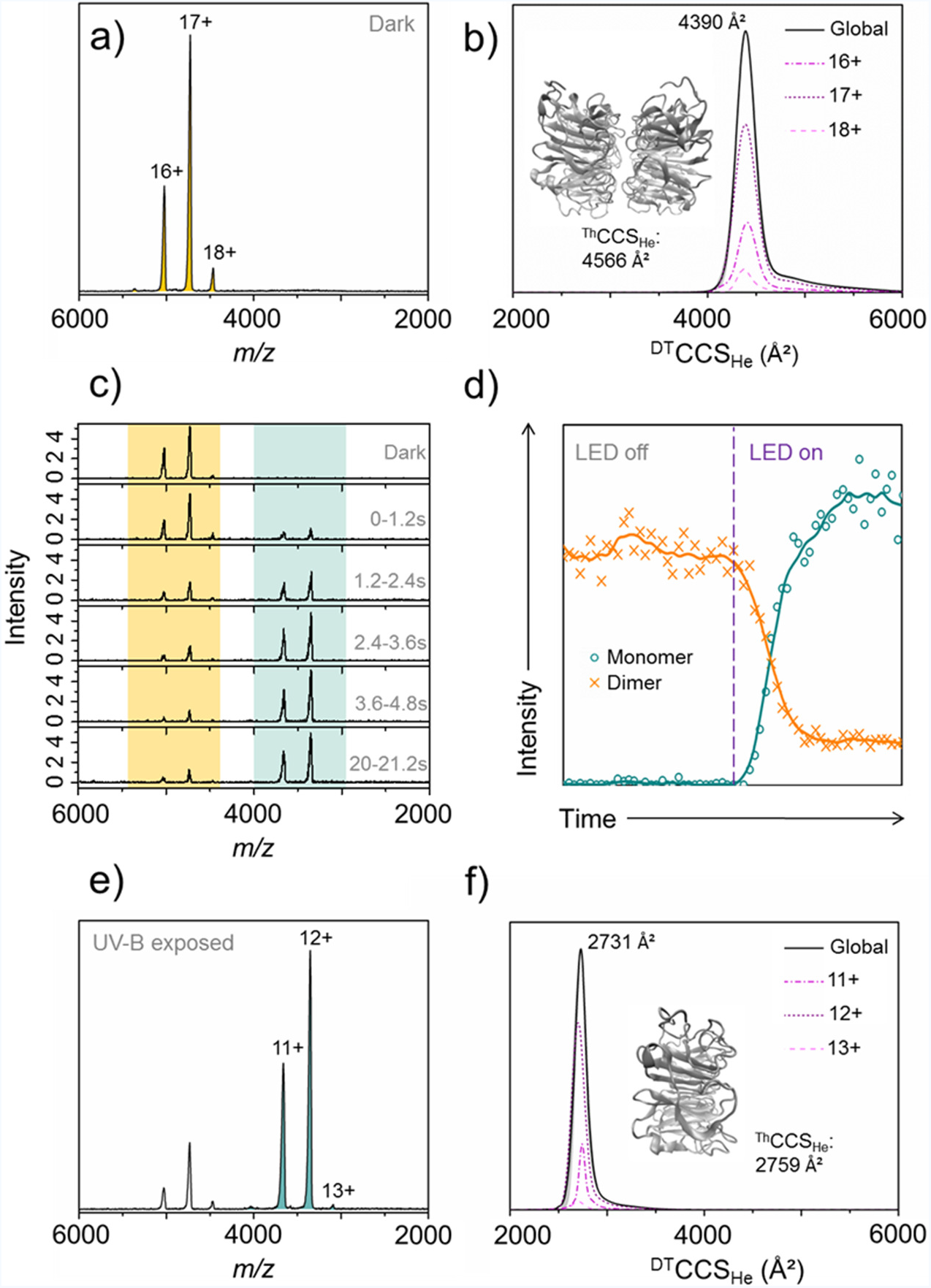
Native mass spectral and ion mobility data of the UVR8 core domain (UVR8^12-381^). The mass spectra (**a, c & e**) and ion chromatogram (**d**) in this figure were acquired using an Ultima Global mass spectrometer, and the ^DT^CCS_He_ (**b & f**) were acquired using a Synapt G2 modified with a linear drift tube. **a)** Native mass spectrum of the UVR8^12-381^ dimer (pale orange peaks) in the absence of UV-B light with charge states labelled. **b)** The monodisperse ^DT^CCS_He_ of the UVR8^12-381^ dimer measured in helium. Inset: energy minimised structure of the UVR8^12-381^ dimer and associated ^Th^CCS_He_. **c)** Mass spectra of UVR8^12-381^ as a function of illumination time using a high-power LED (280 nm, 25 mW, 350 mA). Mass spectra were combined over a period of 1.2 seconds each after the LED was switched on at *t*0. Dimer signal – pale orange; monomer signals – pale green. **d)** Ion chromatogram extracted for the UVR8^12-381^ dimer (orange) and monomer (green). Data to the left of the purple dashed line are from when the ion source tip is in the dark and the data to the right are from when the tip is illuminated. **e)** Native mass spectrum of UVR8^12-381^ following illumination in the source tip with the 280 nm LED for 10 s (to ensure maximum conversion). Residual dimer peaks are still evident, but the spectrum is now dominated by the UVR8^12-381^ monomer (pale green peaks). **f)** The monodisperse ^DT^CCS_He_ of the UVR8^12-381^ monomer measured in helium. Inset: energy minimised structure of the UVR8^12-381^ monomer and associated ^Th^CCS_He_.

When UV-B light from the 280 nm LED illuminates the UVR8^12-381^ dimer in solution throughout data acquisition, the intensity of the UVR8^12-381^ dimer signal decreases with a concomitant appearance and increase in intensity of signal from the UVR8^12-381^ monomer (Figure 3c). The dimer signal decreases in intensity up to 3.6 - 4.8 s of data acquisition, after which it remains constant. The monomer can be seen after up to 1.2 s and increases in intensity up to 3.6 – 4.8 s. The fact that depletion of the UVR8^12-381^ dimer signal is correlated with the appearance of monomer signal is further illustrated by the ion chromatogram for each in Figure 3d. The native mass spectrum of the UVR8^12-381^ monomer is in Figure 3e, which was acquired after the sample was exposed to UV-B in the ion source for 10 s to ensure full conversion. The monomer also presents a narrow charge state distribution (z= 11+ to 13+), which indicates that a single conformational family of UVR8^12-381^ dimers dissociate into a single conformational family of UVR8^12-381^ monomers. This narrow distribution was again confirmed by the ion mobility data acquired in both helium (Figure 3f) and nitrogen (Figure S1b). A global ^DT^CCS_He_ of 2731 ± 5 Å^2^ was obtained from these data, and is in good agreement with the ^Th^CCS_He_ of 2759 ± 32 Å^2^ from MD simulations (Figure S2b) of the energy minimised, published^9^ structure (Figure 3f inset). Much like the UVR8^12-381^ dimer, therefore, the UVR8^12-381^ monomer adopts a well-ordered structure in the absence of its IDRs. The native mass spectra of the UVR8^12-381^ dimer and monomer were measured using three mass analysers (Figures 3 and S3) all revealing comparable results. *In vitro* redimerisation of the UVR8^12-381^ monomer in the absence of UV-B was also measured by native mass spectrometry (Figure S4), occurring over several hours in a manner similar to that observed previously by SDS PAGE.^9^

### Full-length UVR8 exists in numerous conformational families

As we have seen, photoactivation of UVR8^12-381^ results in a straightforward conversion between a single conformational family of structurally ordered dimers to a single conformational family of structurally ordered monomers. In sharp contrast to this, the full-length protein adopts numerous conformational states, both before and after exposure to UV-B. The full-length dimer presents in at least two distributions (Figure 4a), with charge states ranging from z= 17+ to 21+ and from z= 22+ to 31+. Measured ^DT^CCS_He_ values from ion mobility data (Figure 4b and Figure S5a) confirm that these distributions correspond to compact (^~^4500-5400 Å^2^) and more extended forms (^~^5600-7000 Å^2^), respectively. The extended distribution is divided into two sub-populations with the first apex at ^DT^CCS_He_ ^~^6240 Å^2^ and the second at ^DT^CCS_He_ ^~^6660 Å^2^. Because the structure of the core domain appears to be highly stable (*i.e.*, the data of the UVR8^12-381^ dimer in Figure 3) we hypothesised that these populations correspond to different conformational folds of the C- and N-terminal tails, which are thought to be flexible.^9^

**Figure 4.**
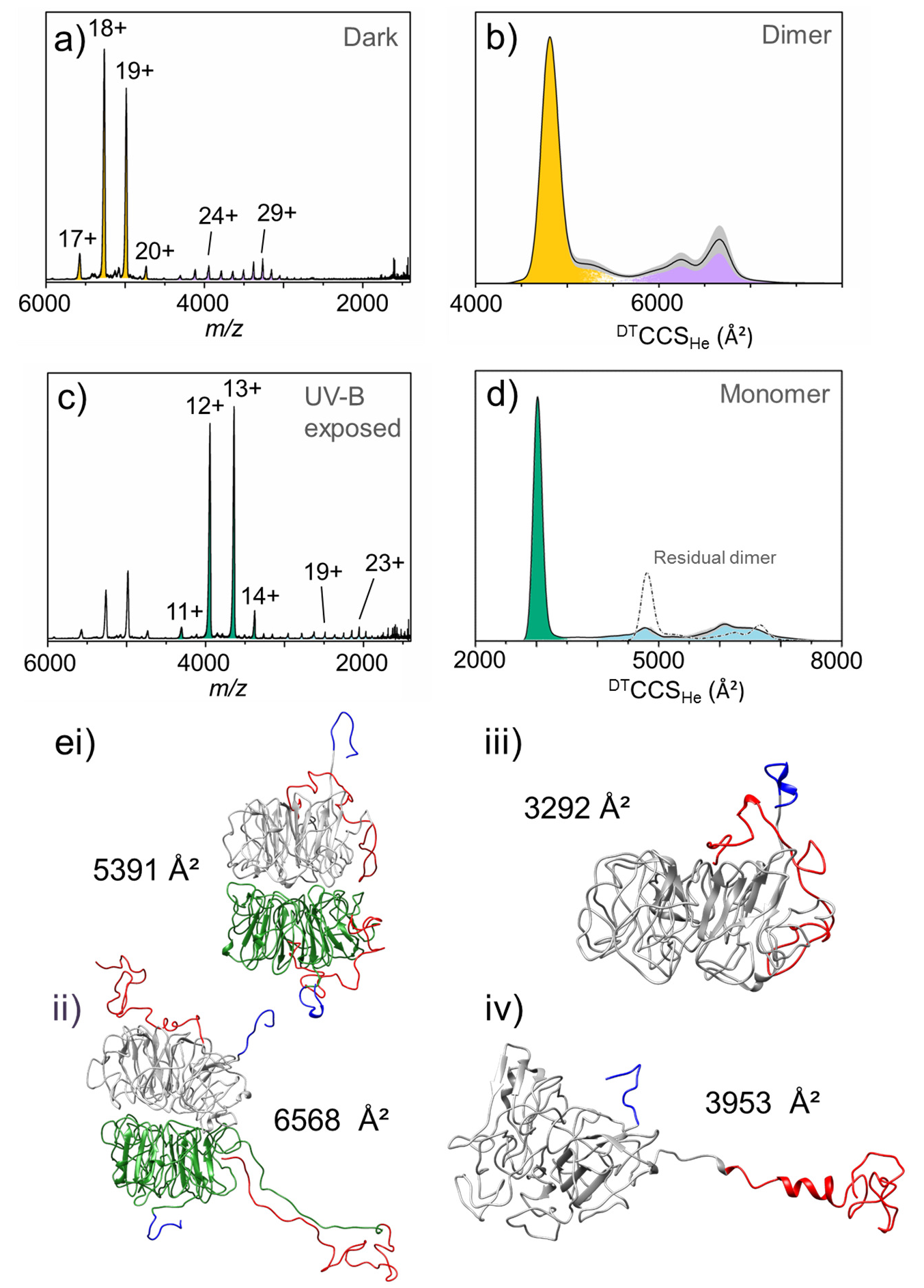
Native mass spectral and ion mobility data of full-length UVR8. All experimental data presented in this figure were acquired using an Agilent 6560 Ion Mobility Q-ToF mass spectrometer. **a)** Native mass spectrum of the full-length UVR8 dimer in the absence of UV-B light with charge states labelled. The full-length UVR8 dimer presents in at least two different conformational families, compact and extended (orange and purple, respectively). **b)** ^DT^CCSD_He_ of the full-length UVR8 dimer measured in helium. ^DT^CCSD_He_ of all charge states were weighted according to the intensity of the peak in the mass spectrum and averaged to get the global distribution. The grey shaded areas represent the standard deviation. **c)** Native mass spectrum of full-length UVR8 following illumination in the source tip with the 280 nm LED for 1 s. Residual dimer peaks are again still evident, but the spectrum is now dominated by the full-length UVR8 monomer, which presents in at least two different conformational families, compact and extended (green and light blue, respectively). **d)** DTCCS_He_ of the full-length UVR8 monomer measured in helium. The residual dimer signal is represented by a dot-dashed line. **e)** Representative structures of full-length UVR8 from gas-phase MD simulations: **i)** compact dimer, **ii)** extended dimer, **iii)** compact monomer, **iv)** extended monomer. The UVR8 core domains are coloured grey and green, the N-termini are in dark blue and the C-termini in red.

To test this hypothesis, we modelled in the C- and N-terminal tails on to the published^9^ structure of UVR8^12-381^ (Figures 1 and S6). In the *in silico* model of the compact dimer, the theoretical collision cross section from MD simulations (Figure S2c) is ^Th^CCS_He_ = 5391± 57 Å^2^ and a representative structure shows both C-termini are close to the β-propeller core of the protein (Figure 4ei). A typical simulated structure (Figure S2d) of the extended dimer, on the other hand, has a ^Th^CCS_He_ of 6568 ± 94 Å^2^ and the C-terminal tails have unravelled into positions well away from the core domain (Figure 4eii). These data strongly suggest that the compact and extended forms of the full-length UVR8 dimer do indeed reflect different conformational folds of the terminal tails, in particular the C-terminus (which is substantially longer than the N-terminus). Consistent with this, lowering the concentration of the sample from 5 μM to ≤ 2.5 μM results in an increase in the proportion of extended species (Figure S7a), and variation in the compact to extended ratio could also be observed upon changing instrumentation (Figure S8). It therefore appears that the different UVR8 dimer conformations can interconvert and that the equilibrium between the states is dependent on the protein environment. The ion mobility data of the full-length dimer, as presented in Figure S5a and Figure 5b, also indicate that – rather than a continuum of conformations between the compact and extended forms as one might expect – a single conformational population dominates the extended form, with a minor population representing a transitional state between it and the compact form.

**Figure 5.**
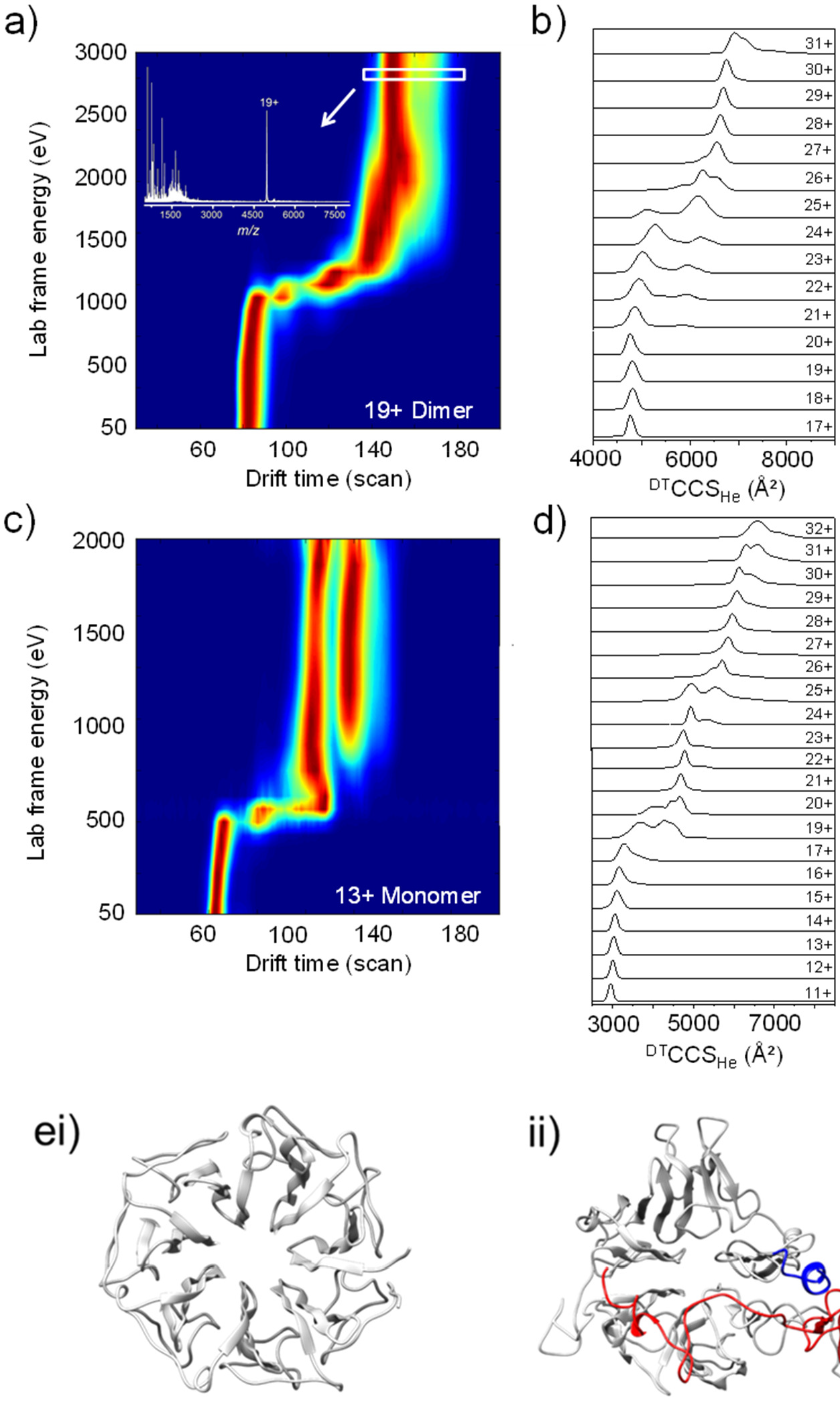
Collisional activation prior to ion mobility (aIMS) and simulated unfolding of full-length UVR8. All aIMS data were acquired using a modified Synapt G2-S mass spectrometer with travelling-wave ion mobility and ion mobility data were acquired using an Agilent 6560 ion mobility enabled Q-ToF mass spectrometer. **a)** aIMS heatmap of the compact 19+ charge state of full-length UVR8 dimer in the gas phase. The drift time of the 19+ charge state was measured as a function of trap collision voltage and converted to lab frame energy (charge × voltage). Inset: mass spectrum recorded at the highest activation energy. **b)** ^DT^CCS_He_ of the full-length UVR8 dimer from ion mobility data. **c)** aIMS heatmap of the compact 13+ charge state of full-length UVR8 monomer in the gas phase. **d)** ^DT^CCS_He_ of the full-length UVR8 monomer from ion mobility data. **e)** Structures of the UVR8^12-381^ (**i**) and (**ii**) full-length UVR8 monomers (both top view) at the end of *in silico* unfolding simulations (Figure S1).

Following exposure of the full-length UVR8 dimer to UV-B light in solution, the mass spectrum becomes dominated by the ion signal of the full-length monomer (Figure 4c). A shorter illumination time (1 s) was chosen than for UVR8^12-381^ in Figure 3 (10 s) to ensure that sufficient dimer remained for its mobility to be measured. The ion mobility data of this dimer were then compared to the non-irradiated dimer (Figure S9); no difference was evident. The monomer also presents in at least two distributions, with charge states ranging from z= 11+ to 14+ and z= 15+ to 32+. Experimental ^DT^CCS_He_ values from ion mobility data suggest the full-length monomer also adopts compact and extended conformations (Figure 4d and Figure S5b). Much like the extended dimer, the z= 15+ to 32+ range for the full-length monomer comprises two sub-populations (with apexes at z= 19+ and z= 23+). Interestingly, the second population spans a very broad ^DT^CCS_He_ range (^~^1000 Å^2^), which suggests the monomer is highly dynamic and can exist in a range of conformational states. Remarkably, this ^DT^CCS_He_ range for the full-length monomer is similar to that encompassing both the compact and extended forms of the dimer (Figure 4d, dot-dashed line) indicating this flexibility is in part dictated by residues that are stabilised in the dimer. MD simulations of the full-length monomer in a compact form (Figure 4eiii) resulted in a ^Th^CCS_He_ value of 3292 ± 39 Å^2^ (Figure S2e), which is in the same region as those measured experimentally (Figure 4d, green). By contrast, the ^Th^CCS_He_ value of the extended monomer (Figure 4eiv & S2f, 3953 ± 63 Å^2^) falls right at the lower limit of the very broad range of experimental values (Figure 4d, light blue). Even fully extending the C-terminal region *in silico* with the core constrained (Figure S10c) leads to a maximal ^Th^CCS_He_ value of 5068 ± 126 Å^2^, which is still less than the ^DT^CCS_He_ range (^~^5600-7000 Å^2^) of the main extended population.

We therefore hypothesised that, as the monomer becomes more extended, it does not simply represent an unfolding of the terminal tails, but of the core protein fold as well. The beginning of this process is evident in the representative structure of the extended monomer following MD simulation in Figure 4eiv). The core domain is unconstrained during these simulations, and is clearly becoming less well-ordered than in the dimer structures (both UVR8^12-381^, Figure 3b inset, and full-length, Figure 4ei&ii), the UVR8^12-381^ monomer structure (Figure 3f inset) and the compact full-length monomer structure (Figure 4eiii). It therefore appears that the homodimeric interface not only maintains the quaternary structure in the dark but also stabilises the tertiary structure of the core domain in each full-length monomer. Because similar unfolding of the core domain is not evident in the monomer where the C- and N-termini have been truncated (*i.e.*, the UVR8^12-381^ monomer, Figure 3e), we hypothesised that the extended IDRs of full-length UVR8 serve to destabilise the β-propeller structure in the absence of the stabilising cross-dimer interactions. Overall, we conclude from these data that the full-length UVR8 structure has access to conformations where one or more of the C- and N-termini are extended (Figures 4eii&iv) and even the core, β-propeller structure can adopt more extended states. The ion mobility data of the full-length monomer, as presented in Figure S5b and Figure 5d, indicate that two conformational populations dominate the extended form, with a third, minor population representing a transitional state between them and the compact form.

### The UVR8 dimer is highly stable in the dark whereas the terminal tails are conformationally disordered

It therefore seems likely that the activation of full-length UVR8 not only requires monomerisation triggered by light but also the sampling of conformational space to enable the C-terminus to mediate protein/protein interactions and thus initiate signalling and regulate UVR8 function. Interestingly, our data indicate that the UVR8 dimer is a highly stable state. To investigate this further, collisional activation prior to ion mobility (aIMS) of full-length UVR8 was conducted in the absence of UV-B light (Figure 5a). Here, the drift time of a mass-selected charge state is measured as a function of trap collision voltage (*i.e.*, activation energy). It can be seen in Figure 5a that the initial compact dimer structure of z = 19+ is stable up to an activation energy of ^~^ 1000 eV, after which there is a transition to longer drift times, which are consistent with a more extended dimer structure. A briefly stable, transitional distribution is evident at ^~^1300 eV which then unfolds further and remains stable up to 3000 eV. The mass spectrum at this energy (Figure 5a inset) reveals that, whereas fragments corresponding to parts of the disordered tails are abundant in the spectrum, no monomer is observed. These data follow a remarkably consistent pattern with the ^DT^CCS_He_ trend measured for the charge states of the full-length UVR8 dimer before photo-activation (Figures 5b and S5a): a compact, stable conformation and a transitional configuration leading to a second stable, extended state. The remarkable resilience of the dimer form in the absence of light was also shown by collision-induced dissociation (CID, see Methods section), with the terminal tails being fragmented under collision while the dimer interface remained intact Figure S11). Altogether, these data confirm the hypothesis supported by the computational data in Figure 4: that the compact and extended conformations of the full-length dimer correspond to different conformational folds of the terminal tails.

### The disordered C-terminus destabilises the core structure in the full-length monomer

To test our hypothesis that the interactions at the homodimeric interface not only maintain the quaternary structure but also stabilise the β-propeller core of each monomer, we conducted aIMS of the compact, full-length monomer (Figure 5c). The initial conformation is only stable up to an activation energy of ^~^500 eV (compared to ^~^1000 eV for the compact dimer) and the subsequent transition is far more abrupt. Moreover, it transitions to drift times that are indicative of highly extended monomer conformations, consistent with the ion mobility data (Figures 4d, S5b and 5d). The aIMS data also suggest that there are two stable conformational families of highly extended monomer, whose drift times remain relatively constant as the lab frame energy is increased from ^~^800 to ^~^2000 eV. This is supported by the native ion mobility data of the full-length monomer in Figure 5d, where the two dominant populations of the extended monomer are present in the ions that present with z = 20+ to 24+ and 26+ to 30 +, respectively. In combination, the aIMS and ion mobility data of the full-length UVR8 monomer strongly suggest that, in the absence of the stabilising cross-dimer interactions, the β-propeller fold of the core domain becomes destabilised when the terminal tails are extended.

It should be noted again that the core of the UVR8^12-381^ monomer, which lacks the IDRs, does not present with such conformational diversity of the core domain (Figure 3). To test our previously stated hypothesis that the IDRs of UVR8, and in particular the longer C-terminus, destabilise the tertiary structure of core domain in the full-length monomer, we conducted simulated unfolding of both the UVR8^12-381^ and full-length monomers (Figure S12). These data confirm that the core domain of the full-length UVR8 monomer is substantially less stable than when the terminal tails are absent (*i.e.*, for the UVR8^12-381^ monomer). This is starkly illustrated by representative structures of the UVR8^12-381^ (Figure 5ei) and full-length (Figure 5eii) monomers following these unfolding simulations. Whereas the UVR8^12-381^ monomer remains well-folded, with little deviation from the β-propeller fold of the published crystal structures^9,10,16^ (Figure 1a), the core domain of the full-length UVR8 monomer has become significantly disordered. From these data we therefore conclude that the extended IDRs act to convert the core fold from the β-propeller structure of the homodimeric state towards highly extended, stable conformations in the UVR8 monomer.

### The extended UVR8 dimer dissociates into monomers more readily than the compact dimer

As we have seen, the UVR8 dimer is a highly stable structure, which fails to dissociate into monomers when in the gas phase and in the absence of UV-B even under unfolding conditions. Indeed, the UVR8 dimer still fails to form monomers in the gas phase under UV-B illumination (Figure S13). It is highly likely, therefore, that water molecules are required for the successful photoactivation of UVR8. The interfacial region of each monomer is highly charged and the dimeric structure is maintained in the dark by cross-dimer salt bridges.^9,10^ A water molecule is thought to be ejected from the interfacial region following photoexcitation of the tryptophan chromophore; it has been proposed that this leads to disruption of a network of H-bonds and the cross-dimer salt bridges.^16^ For the dimers to convert to monomers following photoactivation rather than immediately redimerise, our experiments indicate that additional water molecules that are absent in the gas phase must be required to enter the interfacial region and shield the complementary charges from one another.

Gas phase monomerisation was only possible using surface-induced dissociation (SID), which imparts significant amounts of energy onto the ion in a single, high-energy collision with a functionalised gold surface. A SID voltage of 100 V was sufficient to dissociate the dimer into monomeric units (Figure 6a). SID of compact dimers (*e.g.*, z = 19+ and 20+) results in an asymmetric dissociation pattern in the resulting monomer spectra (Figure 6a orange panels), with many charge states of the monomer evident. By contrast, extended dimers (*e.g.*, z = 25+ and 26+) give a symmetric dissociation pattern in the resulting monomer spectra (Figure 6a purple panels) in which only a narrow range of charge states are produced. The broader charge state distribution in the SID spectra resulting from the compact dimer state results from ‘scrambling’ of charges along the interface. Such scrambling reflects the fact that more protons are transferred to one monomer of the dissociating dimer than to the other, hence an asymmetric charge distribution. Remarkably, what these data reveal is that the extended dimers dissociate into monomers much more readily than do the compact dimers, suggesting additional interactions between the monomers in the compact dimer. Indeed, the representative structures of the compact, full-length dimer (Figure 6b) reveal H-bonds between residues of the C-terminal tail (S396 and K398, which were missing in the published crystal structures)^9,10,16^ with residues on the core domain of the opposite monomer (D367 and E364, respectively). By contrast, the extended dimers dissociate in a far more facile manner, which leads to a lower number of charge states in the dissociated monomer (purple spectra, Figure 6a). SID is thus sensitive to the conformational differences between the compact and extended dimers (Figure S14).

**Figure 6.**
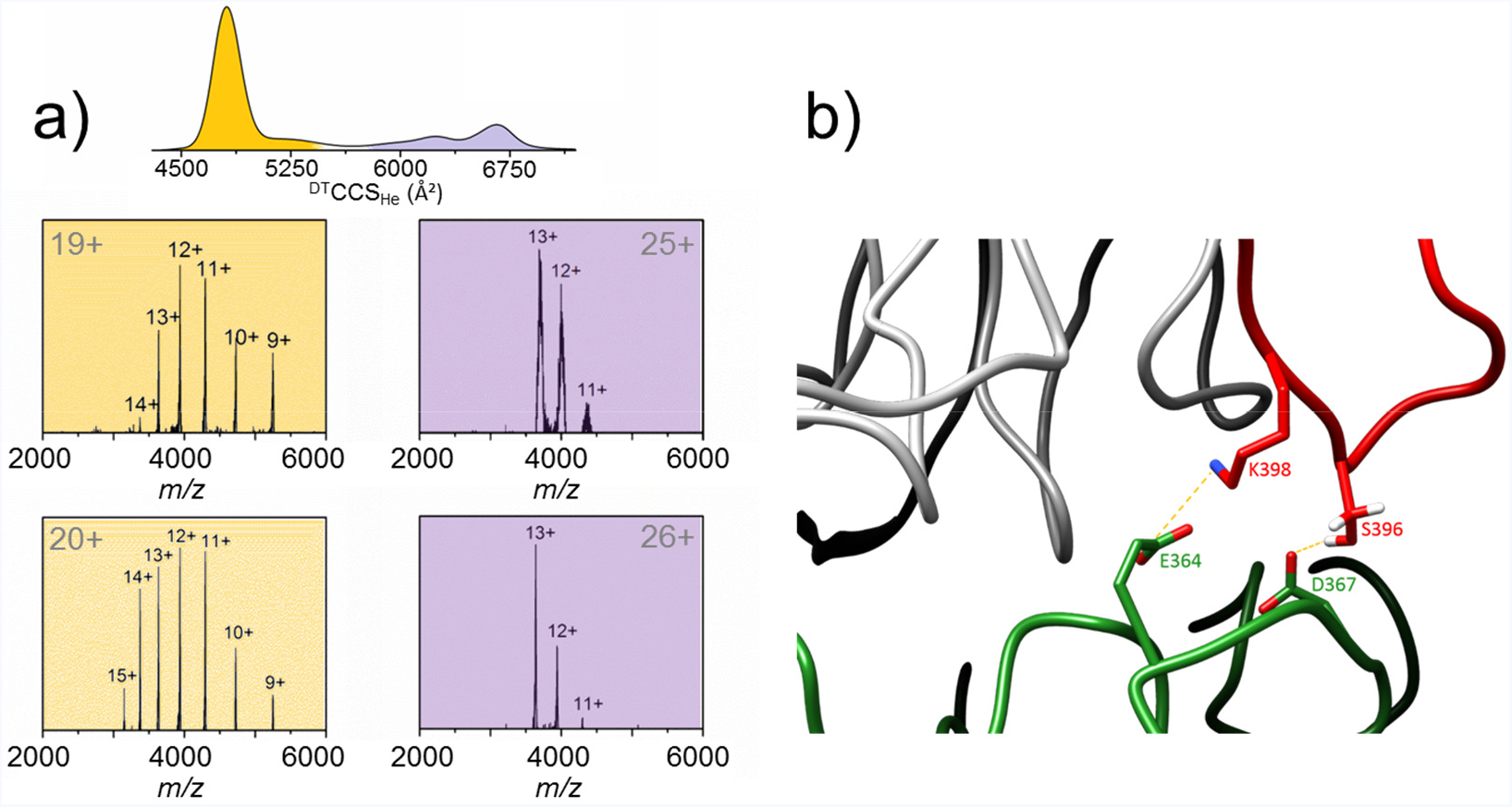
Surface-induced dissociation (SID) of the UVR8 dimer in the gas phase. **a)** Top: ion mobility data showing the compact (orange) and extended (purple) conformations of the full-length UVR8 dimer. Bottom: the 19+ and 20+ charge states of the compact dimer each show an asymmetric dissociation pattern in the resulting monomer spectra (orange panels), whereas the 25+ and 26+ charge states of the extended dimer each show a symmetric dissociation pattern in the resulting monomer spectra (purple panels). **b)** A representative structure of the compact full-length dimer conformation where the C- and N-terminal tails have been modelled on to the published^9^ structure of the UVR8^12-381^ dimer. H-bonds are evident between the C-terminus (red) of one monomer (the rest of which is coloured grey) and the core domain of the opposite monomer (green).

## Discussion

Sunlight is vital to plant life, not only as a source of energy harnessed by photosynthesis, but also as a source of information about their environment. This information is processed by a variety of photoreceptor proteins that absorb light from the UV-B right across the visible spectrum to the far-red.^17^ The plant photoreceptor UVR8 has evolved to respond to and thus protect against the harmful effects of UV light.^7^ UVR8 was the first discovered among only a very small number of photoreceptors^11,18^ known to employ tryptophan residues from the protein backbone as photoactive chromophores. Crystal structures of the core domain where the C- and N-terminal tails had been truncated (UVR8^12-381^)^9,10^ reveal that a pyramid of 4 tryptophan residues comprise the central chromophore, with W233 and W285 being necessary for UV-activation. Photoexcitation of these residues results in disruption of a network of salt bridges that stabilises the homodimeric structure, leading to dissociation into monomers. We have shown that the evolution of the dimer and monomer signals of UVR8^12-381^ under UV-B illumination can be monitored using native ion mobility mass spectrometry. Moreover, these data nicely demonstrate that the decay of the former is correlated with the appearance of the latter, and that a single conformational family of dimers converts to a single conformational family of monomers. In other words, both the UVR8^12-381^ dimer and monomer are well-folded structures.

By contrast, we have revealed that full-length UVR8 possesses considerable conformational diversity. We have identified extended states that are not only accounted for by different conformations of the terminal tails, which were already thought to be IDRs, but also of the β-propeller structure of the core domain, which until now was thought to be relatively rigid. Full-length UVR8 and COP1 interact in the nucleus following UV-B illumination both *in planta*^14^ and in protein extracts.^11^ Although upon UV-B illumination UVR8^12-381^ can monomerise, accumulate in the nucleus^5^ and then interact with COP1^15^, it is missing a highly conserved 27 residue region of the C-terminus (C27) that is required for functional binding. Our data reveal that the full-length UVR8 dimer exists in two main conformational families, one compact and one extended, with a transitional state between them. The compact and extended forms of the dimer are defined by different conformational states of the terminal tails, and in particular the longer C-terminus, and the extended family is more conformationally diverse. Access to extended conformations is consistent with data from size exclusion chromatography, which indicate that the full-length dimer elutes at an apparent molecular mass of ^~^ 150 kDa^10^ despite the calculated mass from its sequence being in the order of 100 kDa. Extended conformations are also supported by previous SAXS analysis, which also suggest the C-terminus is located on the distal side, away from the dimer interface.^9^ Our MD simulations of the compact dimer, however, reveal likely interactions between the C-terminus of one monomer and the core domain of the opposite monomer. These interactions involve residues that were truncated in the published crystal structures.^9,10,16^ Access to both compact and extended conformations, and the fact that we show their relative proportions to be dependent on their environment, provide strong experimental evidence for conformational flexibility of the terminal tails. The fact that we observe two main populations, however, and not a continuum of states, suggests there is an element of order to this flexibility. Consistent with this, previous modelling of the C-terminus suggested that residues 400-421 and 432-440 are likely to be α-helical, with the remaining, unfolded residues predicted to be flexible.^19^

In the absence of UV-illumination, the UVR8 homodimeric structure is very stable. Indeed, we show that the dimer remains intact following CID, whereas the C- and N-terminal tails undergo fragmentation. This is consistent with that fact that un-boiled samples of the isolated protein in solution remain dimeric in the dark even under the denaturing conditions of SDS-PAGE.^9,10^ Remarkably, we further reveal that in the gas phase even UV-B illumination is insufficient to cause significant monomerisation. It is therefore likely that light-activated monomerisation proceeds in two stages. First, photoexcitation in solution triggers disruption of the cross-dimer salt bridges causing initial separation of the monomers. Second, water molecules surrounding the homodimer and within the central, water-filled tunnel^9^ are required to enter the interfacial space in order to shield the salt bridge partners from one another and thus enable the monomers to separate completely. These water molecules will mostly be absent in the gas phase, and are therefore unavailable to stop the salt bridges from reforming. We found that surface-induced dissociation (SID) provided sufficient energy to cause the dimer to dissociate into monomers in the gas phase. SID is equivalent to a substantial increase in temperature, which is consistent with boiling of unilluminated aqueous samples of dimer being necessary to produce monomers on SDS-PAGE.^9,10^ SID also reveals that the extended dimer converts to monomers following SID more readily than the compact dimers do. This may well be owing to the additional interactions evident between the C-terminus of one monomer and the core domain of the other monomer in the compact form (Figure 6b). The extended dimer conformation could therefore represent a state ‘primed’ for activation, leading to a monomeric form with an extended C-terminus that is ready for functional interaction with COP1.

The UVR8 / COP1 interaction is UV-B dependent,^14^ and the dimer with an extended C-terminus is therefore not sufficient for functional binding. Based on this, it is highly likely that both monomerisation and access to the extended conformations of the C-terminus that we observe are required to effect functional binding and therefore signal transduction. Much like for the full-length dimer, and in contrast to the UVR8^12-381^ monomer, the native mass spectrum and accompanying ion mobility data for the full-length monomer reveal numerous conformational states. These are consistent with a single compact form, a transitional state and two highly extended, but nevertheless stable states. In contrast to the full-length dimer, the extended monomeric states cannot be accounted for by extended conformations of the C- and N-termini alone, and also require unfolding of the β-propeller structure of the core domain. Extended monomer conformations are consistent with published *in vivo* data; an additional band appears above the monomer on SDS-PAGE of unboiled protein extracts from *Arabidopsis uvr8-1* expressing GFP-UVR8 variants.^20^ This band increases in intensity with UV-B illumination of GFP-UVR8, is present but unresponsive to UV-B in the constitutive monomer, GFP-UVR8^R286A^ and is not apparently detected for the constitutive dimer, GFP-UVR8^W285F^. All of these trends are consistent with our data.

The existence of highly extended monomeric states suggests that the cross-dimer salt bridges not only serve to stabilise the homodimeric structure in full-length UVR8, but also the β-propeller structure of the core domain. The difference we observe in stability of the β-propeller fold between UVR8^12-381^ (more stable) and full-length (significantly less stable) monomers indicates that the presence of the terminal tails, and in particular the longer C-terminus, serves to destabilise the core fold in the absence of the cross-dimer interactions. Dynamic crystallography of UVR8^12-381^ reported UV minus dark difference densities that not only indicated structural changes around the Trp chromophore and at the dimer interface, but also coordinated, large scale changes, which suggest partial unwinding of the β-propeller fold.^16^ These appear to be driven by motions in blades 5 and 6, which contain the key residues of the chromophore, W285 and W233, respectively. This, alongside our data, suggests that structural rearrangement of the core domain is triggered by light not only to effect monomerisation through disruption of the cross-dimer interactions as a result of motions of W285 and W233, but also to trigger destabilisation of the core fold. Then in the full-length protein the extended C-terminus serves to further force the core fold into the stable, highly extended conformations that are able to interact functionally with COP1.

Why might a highly extended state where the core domain is at least partially unfolded be necessary to function? It has been previously demonstrated that COP1 interacts not only with the C-terminus of UVR8, but also with the UVR8 core domain.^15^ We propose that in order for the rather large COP1 protein (658 residues)^21^ to successfully interact with the UVR8 core domain, the latter must unfold to a highly extended conformation. In support of this are data regarding the W285A variant, which is a constitutive monomer. It interacts functionally with COP1 *in vivo*,^22^ but this interaction is not as strong as that of WT UVR8^11^ because it only does so *via* its C-terminus.^15^ This is consistent with the model proposed above: that one role of photoactivation is to trigger the unwinding of the β-propeller core. Further, one cannot rule out the possibility that full activation of UVR8 is a two photon process, with the second photon absorbed by the monomers to effect a more complete unfolding of the core structure. Consistent with this notion, experiments with a monomeric double mutant of UVR8 (D96N/D107N) show that the monomer is able to absorb UV-B and initiate responses *in vivo*, indicating that dimer photoreception is not essential for UVR8 function.^23^ It is therefore possible that a major role of the homodimeric structure in deactivating UVR8 is to provide the stabilising influence of the cross-dimer interactions over the β-propeller fold in each monomer that we have revealed here. Either way, it appears that the efficacy of the functional interaction between the UVR8 C-terminus and COP1 requires the stabilising interaction between COP1 and the UVR8 core domain, which in turn requires the latter to adopt a highly extended conformation to accommodate the COP1 protein. Unlike COP1, RUP1 and RUP2, which are negative regulators of UVR8, only interact with the UVR8 C-terminus and do so in both its monomeric and dimeric state.^5,24^ Do RUP proteins therefore have a higher affinity for the C-terminus of UVR8, and does the relatively weak affinity of COP1 therefore necessitate its binding to the core domain? This would make sense from a functional point of view because presumably RUP needs to displace COP1 to repress UVR8 function. A higher affinity, in combination with a higher concentration of the RUP proteins resulting from up-regulation by the UVR8 / COP1 complex,^24,25^ would mean they are well placed in terms of competitive binding for the UVR8 C-terminus.

We therefore propose a mechanism in Figure 7a that is consistent with our data and the discussion above. The compact and extended UVR8 dimers represent different conformations of the disordered C- and N-terminal tails and are in equilibrium. Photoexcitation of the tryptophan chromophore disrupts the cross-dimer salt bridges and triggers both monomerisation and unwinding of the core domain. Monomerisation occurs preferentially from the extended dimer and requires water molecules to enter the inter-monomer space to shield salt bridge partners from one another. The C-terminus in the resulting extended monomers is thus both available for functional interaction with COP1 and serves to further destabilise the core domain to yield highly extended conformations. A two-step mechanism has previously been proposed for subsequent interaction between the UVR8 monomer and COP1:^15^ a stabilising interaction between COP1 and the UVR8 core domain followed by interaction between COP1 and UVR8 C-terminus. Here we propose that for the initial stabilising interaction to take hold the UVR8 monomer must adopt a highly extended conformation. The UVR8 / COP1 complex upregulates RUP proteins,^25^ which displace COP1 at the C-terminus through competitive binding. The RUP proteins then help bring the UVR8 monomers back together and mitigate the destabilising influence of C-terminus. This enables the cross-dimer interactions to promote refolding of the β-propeller core. The question has also been asked about whether it is necessary for the RUP proteins to disassociate from UVR8 following reassembly of the homodimer and during subsequent rounds of photoactivation.^15^ Figure 7b offers one possible mechanism whereby docking of COP1 to the extended core domain might facilitate displacement of the RUP proteins should they indeed remain bound.

**Figure 7.**
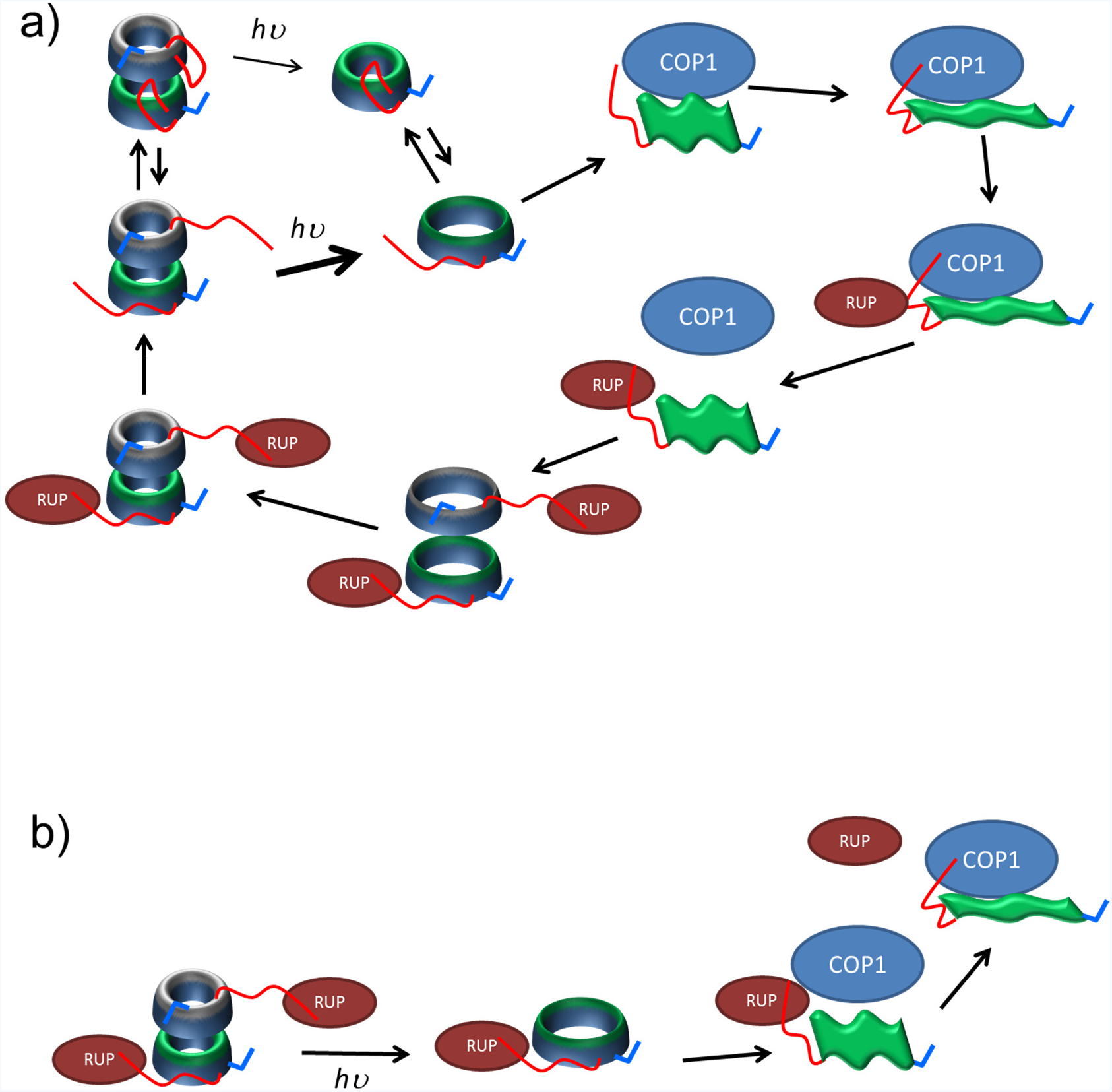
Predictive mechanisms of the protein interactions that mediate UVR8 signalling and regulation. For a full discussion of each mechanism refer to the main text. Consistent with the colour scheme used throughout this article, the full-length UVR8 homodimer is depicted using one green and one grey core domain for the two monomers, each with a red C-terminal and blue N-terminal tail. *hυ* - absorption of a photon. **a)** Cartoon illustrating the proposed mechanism for photodissociation of the compact and extended UVR8 dimers into the various monomer conformations and subsequent interactions between the extended UVR8 monomers (green flag-like structures) and COP1 / RUP proteins. **b)** Possible mechanism for displacement of RUP by COP1 should RUP remain bound to UVR8 during photoactivation.

## Conclusions and Outlook

Organisms throughout nature respond to their ambient light environment through the action of photoreceptor proteins, with functions as various as shade avoidance by plants, neuronal gating and circadian photoentrainment. In most cases, absorption of light results in structural changes in the protein that enable interaction with signalling partners. Such interactions are often mediated by disordered regions. We have demonstrated the power of native mass spectrometry to elucidate the conformational diversity of the plant UV photoreceptor, UVR8. In addition to monitoring the well-described light-triggered changes to quaternary structure, this approach enabled us to detail the different conformations of the flexible N- and C-terminal tails and how they influence the structure of the core domain. Bearing in mind what has been previously reported about UVR8 signalling interactions, it appears that all of these structural changes are likely to be necessary for binding to COP1 and therefore for activation of UVR8 to its fully functional conformation. These data also serve as a good example not only of how IDRs are important for signalling interactions, but also of how the conformation of ostensibly well-folded, highly stable domains can and do change according to functional requirements. We believe this approach will also prove invaluable to future study of the interactions between UVR8 and the COP1/RUP proteins that mediate UV-B signalling and the activation/deactivation dynamics of UVR8. Over recent years, native ion mobility mass spectrometry has emerged as a powerful technique with which to “capture the repertoire of conformational states adopted by protein assemblies”^26^ from a wide variety of biological systems. The experimental approach described here makes the first use of a simple illumination setup that can be adapted straightforwardly to investigate the structural dynamics of any photoreceptor protein. Moreover, the versatility of photoreceptor function means that they are commonly reengineered as optogenetic tools to enable spatiotemporal control over target processes in biology using light. The approach employed here should also prove a powerful addition to the range of techniques with which to investigate, understand and optimise the structural dynamics important to optogenetic function.

## Methods

### Protein expression and purification

The genes that encode full-length UVR8 from *Arabidopsis thaliana* (residues 1-440) and the variant where the C- and N-termini are truncated (leaving residues 12-381, UVR8^12-381^) were cloned into a modified pET28a (Novagen) expression vector using *Nco*I and *Not*I, to provide N-terminal 7xHis and Strep II affinity tags, and expressed as expressed as SUMO fusion proteins as described previously.^9^ Briefly, the plasmids were transformed into *E. coli* Rosetta 2 (DE3) pLysS cells (Novagen), and a single colony inoculated into LB media. Cells were grown overnight at 37°C (pre-inoculum). Terrific Broth was then inoculated with the pre-inoculum, and cells were grown to an OD600 ^~^ 1.0 and induced with 60 μM isopropyl-beta-D-thiogalactopyranoside (IPTG) at 16° C for 20 h. Cells were harvested by centrifugation, snap-frozen in liquid nitrogen, and stored at–80° C. Cells were thawed in buffer A (100 mM Tris-HCl, pH 8.0, 500 mM NaCl, 20 mM imidazole, 1 mM β- mercaptoethanol, 10% glycerol, and protease inhibitors), lysed by incubation with BugBuster reagent for 1 h at room temperature, and the cell debris removed by centrifugation. The supernatant was collected, filtered through a 0.45 μm membrane, incubated in batch with 5 mL Ni-NTA SuperFlow (Qiagen) for 1 h, washed in buffer A, then buffer B (100 mM Tris-HCl, pH 8.0, 500 mM NaCl, 10 mM ATP, 2.5 mM MgCl2, 1 mM β- mercaptoethanol, protease inhibitors), and lastly with buffer C (100 mM Tris-HCl, pH 8.0, 150 mM NaCl, 1 mM β-mercaptoethanol, protease inhibitors). Protein was eluted with buffer C supplemented with 300 mM imidazole. Eluted protein was incubated in batch with 1 mL Strep-Tactin XT Superflow (IBA-lifesciences) for 1 h and washed with buffer C. UVR8 was cleaved and eluted from the resin by incubation with SUMO protease at 4° C for 16 h, and concentrated to 10 mg/mL using a Vivaspin 2 (10K MWCO, Sartorius). UVR8 was then dialysed to 250 mM ammonium acetate, pH 6.8, using a Slide-A-Lyzer Dialysis Cassette (10K MWCO, ThermoFisher Scientific).

### Mass Spectrometry and irradiation experiments

Mass spectrometry experiments were performed on a modified traveling-wave ion mobility enabled Synapt G2-S (Waters, UK) described previously,^27^ a Ultima Global (Micromass, UK) optimised for high mass range, an Agilent 6560 Ion Mobility Q-TOF (Agilent, California) and a modified Synapt G2 with a linear drift tube in place of the standard triwave assembly. nESI tips were pulled in house from borosilicate glass capillaries on a Flaming/Brown Micropipette Puller (Model P-97, Sutter Instrument Co., USA). The sources of the Synapt and Ultima instrument were modified to accommodate a high-powered 280 nm LED (Thorlabs), focused on the nESI tip (Figure 2) allowing irradiation of the sample solution before it enters the mass spectrometer. 5-20 μM UVR8 full-length and core were measured before, during and after irradiation. Data were analysed using MassLynx v4.1 (Waters Corporation, USA), OriginPro 9.1 (OriginLab Corporation, USA) and Microsoft Excel 2010 (Microsoft, USA).

### Native Mass Spectrometry

Native mass spectra were recorded on an Ultima Global (Micromass, UK) employing gentle source conditions with a capillary voltage of ~1.4 kV, cone voltage of 35 V and all radio frequencies set to 0.

### Ion mobility Mass Spectrometry – CCS measurements

Collision cross section (CCS) measurements were performed on an in-house modified Synapt G2 mass spectrometer. The original traveling-wave ion mobility assembly of the instrument was replaced with a linear drift tube with a length of 25.05 cm. Measurements were made in helium at 298.15 K and a pressure of 1.98- 2.01 Torr was kept constant throughout each run. Spraying conditions were set to be as gentle as possible with an applied capillary voltage ranging from 1.1-1.3 kV, a sample cone voltage of 15 V and source temperature of 298.15 K. Drift times were recorded at a minimum of six voltages ranging from 220-120 V. All experiments were conducted in triplicate over at least two separate days. The drift times obtained were converted into rotationally averaged ^DT^CCS_He_ values using the Mason-Schamp equation.^28^

Additional CCS measurements were performed in nitrogen on an Agilent 6560 Ion Mobility Q-TOF. VCap was set to 1.8 kV with a fragmentor voltage of 500 V. The drying gas was down-regulated manually to 250 μL min^−1^ and drift times were recorded over five voltages ranging from 1700-900 V. Measurements were made in triplicate over at least two separate days. Drift times were converted into ^DT^CCS_N2_ using the Mason-Schamp equation. ^DT^CCS_N2_ obtained were further converted into a single, global ^DT^CCS_N2_ for each species by summing the ^DT^CCS_N2_ of all measured charge states weighted according to their individual intensity in the mass spectrum.

### Collision-induced dissociation (CID) and collisional activation prior to ion mobility (aIMS)

To follow aIMS, the species of interest was mass selected in the quadrupole of a Synapt G2-S and its drift time measured as a function of trap collision voltage which was increased incrementally from 4-150 V. Fragmentation at a selected collision voltage (110-130 V) was analysed and compared between species. Heatmaps were plot using CIUsuite.^29^

### Gas phase illumination

The gas phase illumination experiments were conducted using a modified TWIMS-enabled Q-ToF mass spectrometer that allows UV photodissociation (UVPD) within the instrument, by overlapping of a laser beam with the ion beam within the transfer cell region.^27^ The instrument modification includes an ion gating and trapping procedure that allows ions to be stored for several seconds, enabling UVPD. Illumination was achieved from the fourth harmonic (266 nm) of Continuum Minilite II Nd:YAG laser at a repetition rate of 10 Hz. Typical times used in these experiments were: 1 second filling and 1 second trapping. During these periods, the laser was firing and ions experienced between 10 and 20 laser shots. The energies used range between 0.4 and 1 mJ per pulse. The overall transmission of the energy between the output of the laser and the entrance of the transfer cell was measured as 50%. As a control, equivalent experiments were conducted in the solution phase by placing the sample solution in the path of the 266 nm laser beam outside of the mass spectrometer. This was to confirm that UVR8 dissociates into monomers under similar conditions but in the solution phase (Figure S13d).

### Surface-induced dissociation (SID)

The trap cell of the Synapt G2-S instrument was replaced with a shortened version and a SID device inserted between the shortened trap cell and ion mobility cell. Voltages were optimised to maximise transmission through the device. The species of interest were mass selected in the quadrupole and directed towards a gold surface where they experience a single high energy collision. Fragments were then refocused and guided through the rest of the instrument for analysis.

### Constructing an initial model of full-length UVR8

To model the N- and C-terminal loops missing from the crystal structure, I-TASSER^30-33^ was used to create models for each of the two chains from PDB 4dnw, using 4dnw as a template. The models were then aligned with the crystal structures, and the coordinates of the modelled loops added to the crystal structure. Using the loops from the top-scoring I-TASSER models (as ranked by I-TASSER’s C-score) for each chain leads to clashes between the C-terminal loops since these loops were built extending into the dimer interface (Figure S6a-b). Therefore, the model with the second-highest score for each was used (Figure S6c). Of course there is significant uncertainty as to the actual conformations of the terminal loops. The purpose of the modelling carried out in this study, however, is not to generate definitive conformations of the compact and extended forms of the protein, which would require far more extensive sampling than carried out here, but rather to demonstrate the effect of changes in loop conformations on CCS values by generating reasonable examples of the compact and extended forms. Crucially, for both the full-length monomer and dimer, two methods were used to generate an extended form – using solvated and gas-phase dynamics from different starting points – which resulted in similar ^Th^CCS_He_ values (Figure S2 and Figure S10).

### MD simulations of solvated protein

Molecular dynamics (MD) simulations of the solvated protein were run using single-precision Gromacs 5.0.4^34^ with the Amber 14 force field.^35^ All simulations used a 10 Å cut-off for electrostatic and van der Waals interactions, the lincs algorithm^36^ for bond constraints, a 2 fs time-step, Particle Mesh Ewald electrostatics and periodic boundary conditions. The system setup was as follows: (i) the protein was placed in a solvation box of minimum 12 Å around the protein, and counter-ions were added; (ii) energy minimisation was carried out and the system was thermalised at 300 K for 100 ps with constant pressure and 100 ps with pressure coupling (using the Parrinello-Rahman barostat), with constraints on the protein. All subsequent MD simulations in solvent were carried out using pressure coupling. First, 50 ns of conventional MD simulations were run, followed by 20 ns of simulated annealing to encourage additional conformational sampling, with the following 1.25 ns cycles: 1 ns at 300 K,50 ps of heating to 350 K and 150 ps at 350 K and 50 ps cooling to 300 K.

### Calculating the Theoretical CCS (^Th^CCS_He_)

To obtain structures for CCS calculations, gas phase simulations were run at 310 K. These were run using double-precision Gromacs 4.6.1^34^ using twin range cut-offs for electrostatic interactions, infinite coulombic and van der Waals cut-offs (technically 999.0 nm) and a 2 fs time-step. The setup consisted of energy minimization followed by 50 ps of thermalisation at 250, 280, 290 and 300 K. For the UVR8^12-381^ monomer and dimer, the crystal structure (PDB 4dnw) was used as the starting point. For the full-length monomer and dimer, the starting point was the structure after 20 ns simulated annealing. CCS values were calculated using EHSSrot.^37^ every 500 ps, and the ^Th^CCS_He_ values are the averages from the last 10 ns of these simulations.

### Generating extended conformations

An initial extended conformation of the dimeric protein was created by pulling the centre of mass of the α- carbons of the C-terminus (residues 378-440) from the centre of mass of the α-carbons of the core of each monomer at a rate of 0.01 Å per ns for 10 ns (Figure S6d). This structure was then relaxed for 20 ns using the same annealing procedure as described above for the compact dimer, which was sufficient for convergence of the radius of gyration (Figure S6e). The resulting structure was taken as a starting point for generating gas-phase structures for calculating CCS values for the dimer as well as the monomer. To test the effect of maximally extending the C-terminal region of the dimer and monomer, the C-terminus was pulled during gas phase simulations (since the water box would be prohibitively large for the fully extended state) at a rate of 0.02 Å per ns for 10 ns (Figure S10a,c). Here, position constraints were applied to the core structure, as otherwise partial unfolding of the protein was observed (presumably, unfolding the C-terminus in the gas phase requires significantly more energy than in solution). The protein was then allowed to relax with unrestrained MD simulations Figure S10b,d) to generate additional extended conformations.

### Unfolding simulations

Unfolding simulations were performed to qualitatively assess the relative stability of the UVR8^12-381^ and extended, full-length monomers, starting from the structures and velocities after 10 ns of 300 K MD. Simulated annealing simulations were performed with the temperature cycled between T_1_ and T_2_ with a periodicity of 1 ns: 400 ps at T_1_, 100 ps heating to T_2_, 400 ps at T_2_ and 100 ps cooling to T_1_ with the following T_1_-T_2_ values: 300-400 K, 350-425 K, 350-450 K and 350-500 K. The RMSD for the UVR8^12-381^ residues for each structure (Figure S12) suggests that the full-length monomer is significantly less stable, as the RMSD for the core monomer does not increase significantly even after 10 ns of the harshest annealing. Figure 5e shows each structure at the end of this annealing simulation.

## Acknowledgments

The authors thank Dr. Roger J. Kutta for help with the early stages of instrumentation development. ARJ thanks The University of Manchester and the NMS from the UK Government Department for BEIS for funding. ISC was supported by a PhD studentship from the UK EPSRC. AT is supported by BBSRC grants BB/L002655/1 and BB/L016486/1, which is a studentship award with additional financial support from Waters Corp. LAD-R is supported by a PhD studentship from Consejo Nacional de Ciencia y Tecnología (CONACYT). GIJ thanks the University of Glasgow for the support of his research. The authors would like to acknowledge the assistance given by IT Services and the use of the Computational Shared Facility at The University of Manchester.

## Author Contributions

AT and ISC contributed equally to this work. All experimental work was conducted by ISC, AT, LAD-R and BB, and computational work was conducted by LOJ. Data were analysed and interpreted by all of the authors. The study was conceived and designed by ARJ, PB and BB with GIJ and JMC. ARJ wrote the paper with input from all of the authors.

**Supporting information** is available.

